# Post-embryonic tail development through molting of the freshwater shrimp *Neocaridina denticulata*

**DOI:** 10.1101/2024.03.13.583832

**Authors:** Haruhiko Adachi, Nobuko Moritoki, Tomoko Shindo, Kazuharu Arakawa

## Abstract

**Background:** Understanding postembryonic morphogenesis through molting in arthropods has recently become a focus of developmental biology. The hierarchical mechanisms of epithelial sheet folds play a significant role in this process. *Drosophila* is a well-studied model for holometabolous insects, with extensive research on imaginal disc growth. While developmental processes in other arthropods have been described, live imaging of morphological changes is challenging due to the macroscopic movements and hard cuticles. *Neocaridina denticulata*, a crustacean, presents unique tail morphogenesis through molting, which makes it the potential model. This study investigated the development of the tail in *Neocaridina denticulata* through histological analysis and *in vivo* live imaging using fluorescent probes. This study also performed long-read sequencing of the whole genome for future genetic tools.

**Results:** The tail of *Neocaridina* was found to undergo two major changes with the first ecdysis. Firstly, the branches of the uropods are cleared, and secondly, the telson undergoes convergent elongation. Cross-sectional analysis revealed that uropod and telson branching occurs immediately after hatching in the form of cuticle branching. The surface structure of the developmental tail suggested that telson elongation is achieved by the extension of anisotropic furrows in the cuticle during ecdysis. Anisotropy of cuticle furrows was associated with the epithelial cell shape, and the anisotropy of cell shape was found to occur during development from post-hatching. We also established an *in vivo* live imaging system with UV-LED resin and detected the changes of tail development over time. *in vivo* live imaging analysis revealed that telson contraction occurs gradually prior to ecdysis. Furthermore, we have also provided a draft genome of *Neocaridina*.

**Conclusion:** *Neocaridina denticulata* is a valuable model for studying morphogenesis in arthropods through molting. The tail undergoes complex changes involving cuticle branching, anisotropic furrows, and cellular dynamics. *in vivo* live imaging system provides insights into the developmental process, and the draft genome enhances the potential for genetic tools in future studies. This research contributes to the understanding of arthropod morphogenesis and provides a foundation for further developmental and cytological investigations in *Neocaridina*.

## Background

Understanding the morphogenesis of organisms is a crucial issue in developmental biology. Recently, attention has shifted towards not only embryogenesis but also post-embryonic morphogenesis, particularly in arthropods through molting (1). Among arthropods, insects are particularly well studied for this purpose. Insects can be classified into different modes of development, such as holometabolous, hemimetabolous, and ametabolous insects (2, 3). The developmental process of metamorphosing holometabolous beetles and hemimetabolous treehoppers, respectively, involves universal hierarchical mechanisms. That is the unfolding of epithelial sheet folds and the formation of the folds of different scales. (1, 4-7). In *Drosophila*, a model organism for holometabolous insects, the growth of imaginal discs has been extensively studied from molecular genetic to mechanistic perspectives (8, 9). During the prepupal-pupal period, the individuals are macroscopically immobile while the morphogenetic process occurs. Therefore, *in vivo* live imaging studies are being performed in *Drosophila* (10-12). On the other hand, in hemimetabolous insects, particularly *Gryllus*, it is becoming possible to describe the developmental process and perform some advanced genetic manipulation (13, 14), however live imaging of morphological changes through molting is not yet available. This is because individuals move macroscopically until just before ecdysis, and are covered by a hard, colored cuticle. It is important to study how morphogenesis is achieved in individuals that remain in a state of macromotion until just before ecdysis. Recently, there has also been a growing focus on macro-movement and morphogenesis, it has been found that muscle-mediated macromotion plays a crucial role in morphogenesis in the cnidarian *Nematostella vectensis* (15). Among arthropods, crustaceans move macroscopically just before ecdysis, similar to hemimetabolous insects. Some species in crustaceans also induce morphological changes through molting. Morphogenesis in crustaceans through molting has also not been well analyzed. In this study, we focused on *Neocaridina denticulata*, which is one of the most transparent species.

Genetic tools are currently being developed also for *Daphnia magna* as a research model for crustaceans (16-18). The marine amphipod *Parhyale* has also been recently used as a model organism for crustacean developmental and cellular phenomena (19). The basic genetic tools have also been established (20). Additionally, an amazing live imaging technique using adhesion has been established in *Parhylae* (21) and was used in the comparative analysis of leg regeneration and developmental phenomena (22, 23). On the other hand, *Daphnia* and *Parhyale* have almost the same morphology as their parents from birth, with few tissues undergoing extreme morphogenesis through ecdysis. In contrast, *Neocaridina* exhibits extreme changes of tail morphology through the first molt (24, 25). In *Neocaridina*, the developmental process of the body and the digestive system has been described (24-26). Observational studies have also been conducted on muscle and nerve development during tail development using fixed specimens (27, 28). However, the details of the tail morphogenesis through molting, including the short time scale dynamics, have not been fully clarified. A recent review has described *Neocaridina denticulata* as a potential model organism for decapods due to its ease of husbandry and reproduction (29). Recently, CRISPR/Cas9 knockout experiments have also been conducted with transcriptome data in the congeneric *Neocaridina heteropoda* (30). However, there are currently no publicly available genome assemblies for *Neocaridina*, which could be used to develop genetic tools in the future.

In this study, we investigated tail development in *Neocaridina denticulata* through histological analysis and *in vivo* live imaging using fluorescent probes. Additionally, draft genomes were obtained using nanopore long-read sequencing to establish future genetic tools. This study contributes to a comprehensive understanding of the mechanisms of morphogenesis through molting in arthropods and future developmental and cytological studies in *Neocaridina*.

## Results and Discussion

### The changes of the tail shape through first ecdysis

The structure of a shrimp tail comprises a telson and four uropods. *Neocaridina* undergoes significant changes in the uropod during the first ecdysis after hatching (24, 25). Also in this study, scanning electron microscope (SEM) analysis confirmed that the branching structure corresponding to the uropods was not observed immediately after hatching, and that the branch appeared after ecdysis (Fig. 1a). Furthermore, we recorded the ecdysis process in time-lapse and observed that the change in tail shape occurs within a few minutes (Fig. 1b and movies 1 and 2). The time-lapse imaging also revealed the development of the tail primordium with some level of detail. Although the presence of uropod primordia was unclear in the early stage, the branching of the primordia of uropods and telson became evident in the late stage (Fig. 1b and Mov. 1 and 2). In a millipede, *Niponia nodulosa*, appendages also develop through molting, and it has been reported that they develop inside the transparent protrusion on the surface of the body segments (31). No such structure protruding was observed in the case of *Neocaridina* (Fig. 1a). This could be due to structural differences between regular legs and uropods.

**Fig. 1.**
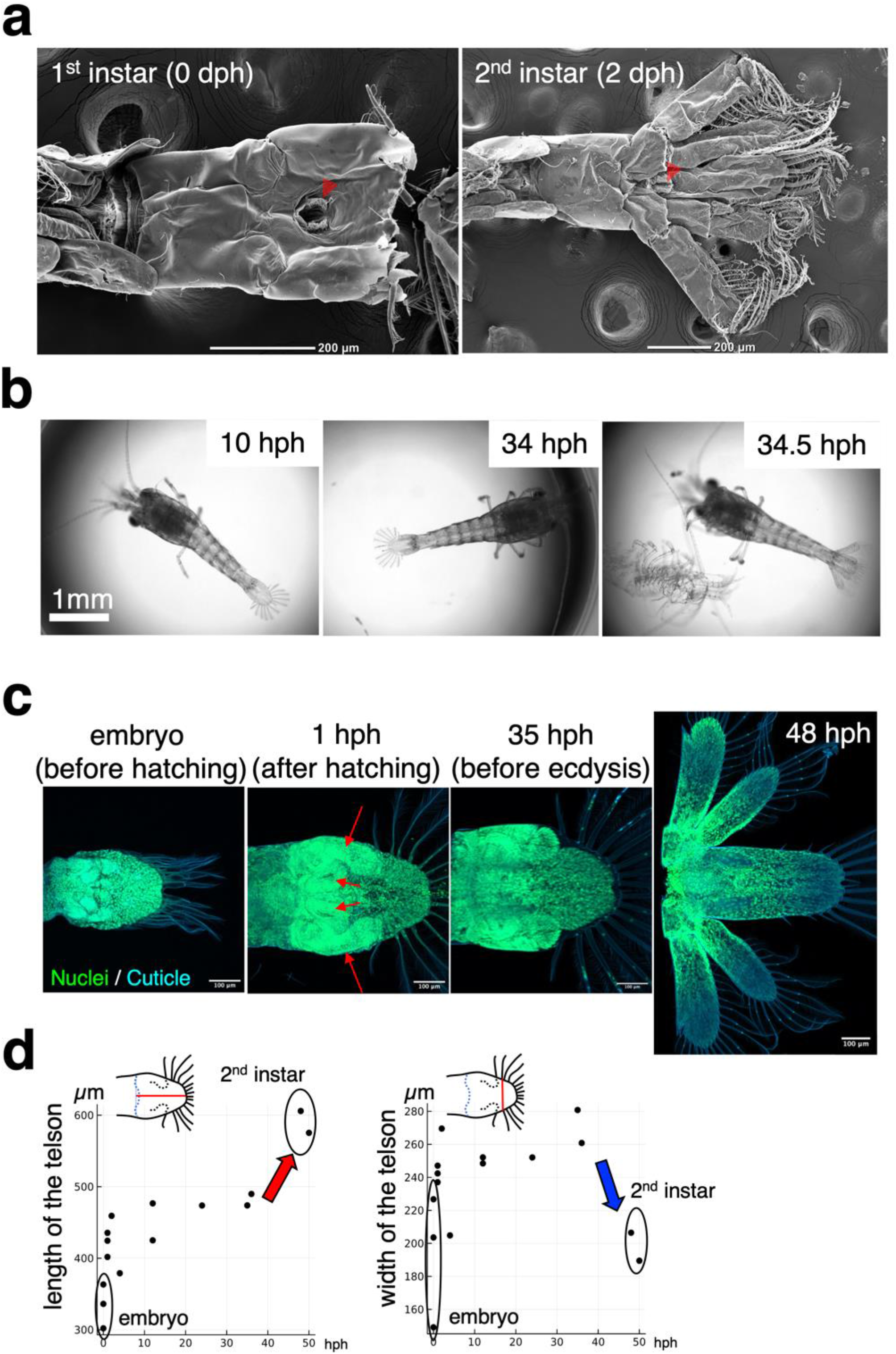
Morphogenesis of the tail of *Neocaridina denticulata*. (a) SEM image of the tail of *Neocaridina denticulate*, 1st instar (0 days post-hatch) and 2nd instar (2 days post-hatch), taken from the ventral side. The red arrowhead indicates the anus. (b) Time-series images of molting. Extracts from the videos at 10 hours post-hatching (10 hph), 34 hours post-hatching (34 hph), and 34.5 hours post-hatching (34.5 hph) are included. (c) The developmental tail was imaged using fluorescence. Nuclei were stained with SYTO13 and the cuticle was detected through autofluorescence upon exposure to a 405 nm wavelength laser. (d) Morphological changes over time were analyzed. The length of the specimen was measured from the root of the uropod to the tip of the telson, while the width was measured between the second setae on the proximal side.

Changes in tail shape during development were investigated using confocal microscopy. Nuclear staining and autofluorescence of the cuticle upon exposure to a 405 nm wavelength laser were used for visualization. The tail of the embryonic period was examined in addition to the time series since hatching. At around one week after incubation, the embryo begins to form an eye and make a beat. The tail was taken out by dissecting the embryo of that period. Upon hatching, bifurcated structures, namely the uropod primordia and the telson primordia, were observed (Fig. 1c). The size of the telsons and telson primordia remained relatively constant between hatching and ecdysis, with notable changes occurring from embryo to hatching and through the first ecdysis. From the embryo stage to hatching, both the length and the width of the telson increased (Fig. 1d). On the other hand, through the first ecdysis, the length increased while the width decreased (Fig. 1d).

The initial molt results in two significant alterations to the tail. Firstly, the branches of the uropods are cleared, and secondly, the telson undergoes convergent elongation.

### Time-series changes in the cross-sectional structure of the tail

The following analysis focused on changes in the cross-sectional structure of the tail. Toluidine blue staining was used to examine the cross-section of the tail during the development process (Fig. 2a). The results showed that just before ecdysis, the uropod primordia and the telson primordia were completely separated. Shortly before ecdysis, the interstitial ECM structure of the toluidine blue-red hyaline was revealed (Fig. 2a). It is believed that the structure may play a crucial role in causing morphogenesis through the first ecdysis within a few minutes, as it is not present immediately after hatching (Fig. 2a). Further research is required to identify this molecule in the future. To investigate the boundary between the uropod primordia and the telson primordia immediately after hatching, we analyzed the microstructure using transmission electron microscopy (TEM). Immediately after hatching, the uropod primordia and telson primordia were found to be isolated by the cuticle (Fig. 2b). In other words, the exoskeleton’s cuticle is already branched immediately after hatching. To confirm this, calcofluor white staining was performed, which stains chitin and cellulose. As a result, consistent with the results of TEM analysis, it was found that the cuticle containing chitin branched so as to isolate the uropod primordia and the telson primordia (Fig. 2c). These findings suggest that the uropod and telson primordia are closely connected from the moment of hatching, enclosed within the cuticle, and that the separation occurs during development (Fig. 2d).

**Fig. 2.**
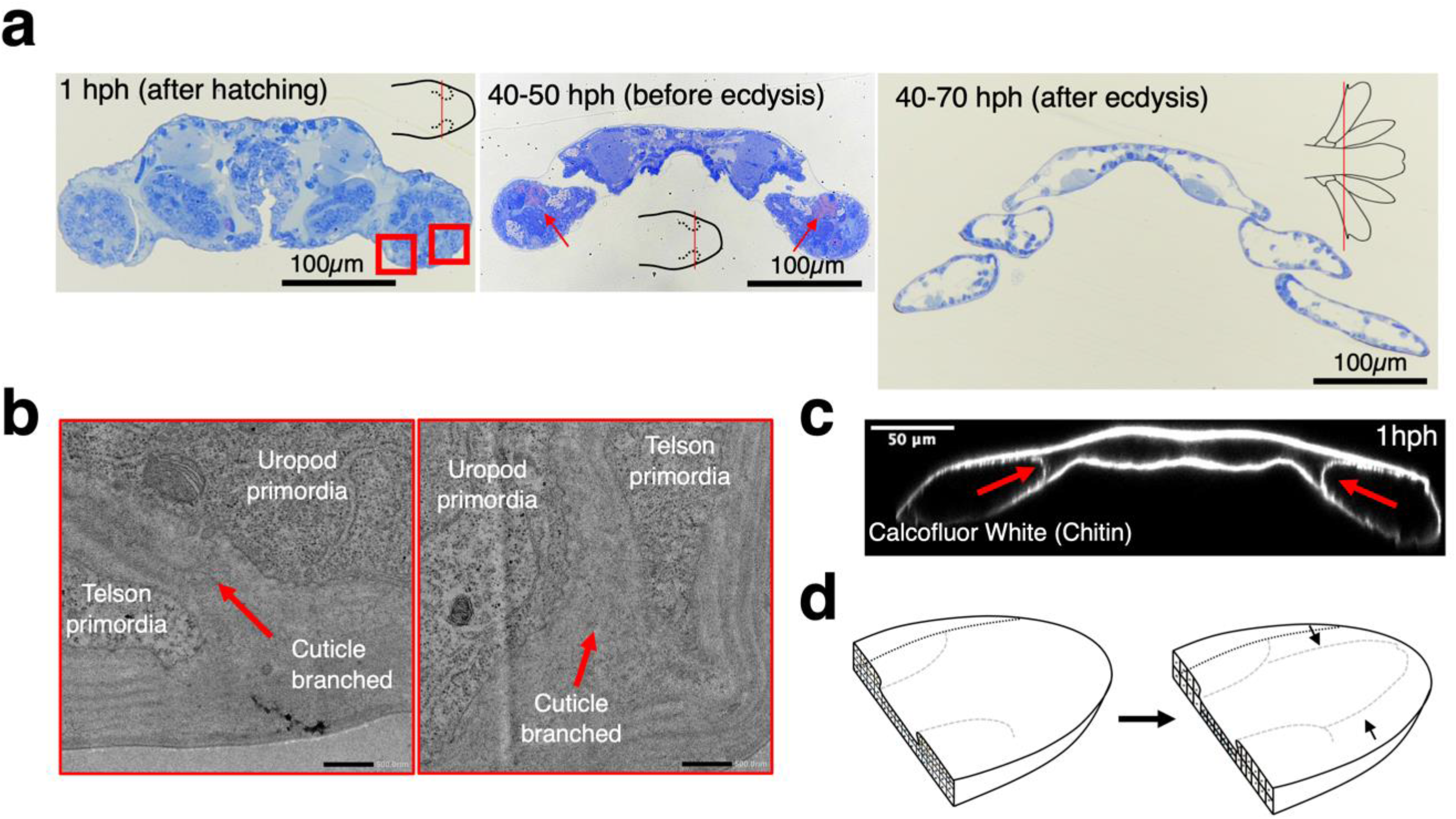
Cross-sectional structure of the developmental tail. (a) Toluidine blue staining of semi-thin section. The plane of section in 1st instar larvae is exactly where the uropod primordium is present. The red rectangles indicate the enlarged areas in (b). The red arrows show toluidine blue-red-purple positive sites, which may indicate the presence of mucopolysaccharides. (b) Transmission electron microscopy (TEM) image from (a) red rectangle regions in 1hph. The red arrows indicate the cuticular branch separating the uropod and telson primordia. (c) Optical section of calcofluor white staining tail in 1 hph. The red arrows show the cuticular branch separating the uropod and telson primordia. (d) Scheme of the cellular structure of the tail primordium during the molting process. Already in the early stages of development, the uropod and the telson primordium are present in an overlapping state (left), although separated by the cuticle, and the overlap is resolved during development (right).

Previous studies have suggested that the development of epidermal and neural tissue in some crustacean larvae is less susceptible to heterochrony than the development of mesodermal muscles (27, 28). Our results show that the cuticle, as well as the epidermal cells, are well-developed in the early stage. This means that just prior to hatching, the patterning of the epithelium of uropod is completed. To further investigate uropod formation, conventional analysis of embryonic development could be examined to clarify the developmental mechanism. The expression of the Hox gene during embryogenesis may be related to this, as observed in the appendages of other crustaceans (32, 33). On the other hand, a fascinating nonconventional phenomenon is the branching of the cuticle. It is still unclear how the epithelium derived from the uropod primordium and the epithelium derived from the telson primordium cooperate to achieve such a structure. Thus, further studies are needed in the future.

### The developmental process of primordium surface structure

The analysis above has enhanced our comprehension of the formation of the branching structures of the uropod and telson primordia. However, our understanding of the convergent elongation of the telson during ecdysis remains inadequate. Since the tail morphogenesis through ecdysis is a brief period of time, the existence of morphogenesis by the folding and unfolding system, as confirmed in insects (1), is expected. This means that the folding structure of the primordia just before ecdysis should allow such a short transformation. When the primordial surface just before ecdysis was exposed by dissection and SEM analysis was performed, anisotropic furrows were confirmed on the surface of the primordia (Fig. 3a). In contrast, no such structures were found on the surface after ecdysis (Fig. 1a). Similar to the morphogenesis through ecdysis in some insects, the unfolding of these furrows is thought to allow a short period of ecdysis transformation, especially elongation. The same analysis was performed on primordia at mid-development and confirmed the cell-like shape corresponding to the direction of the furrows (Fig. 3a). SEM analysis can produce microstructural artefacts due to the ethanol dehydration and freeze-drying processes. Therefore, the microstructure of the cuticle was also observed using calcofluor white staining with confocal microscopy. As in the SEM analysis, anisotropic furrow-like structures were observed in the primordia just before ecdysis (Fig. 3c). No such structures were present jsut after hatching, suggesting that they formed during post-hatching development (Fig. 3c). Next, phalloidin staining was performed to confirm cell-like structures detected by SEM analysis at mid-stage. Samples taken just after hatching and just before ecdysis could not be successfully stained with phalloidin for the outline of the epithelial cells, although the samples could be seen F-actin in neurons, muscles and some mesenchymal-like cells. This may be due to the cuticle, as the outline of the epithelial cells could be clearly seen when the cuticle was detached by apolysis, possibly before the formation of a new cuticle. The cell shape of the uropod primordium at 30 hours post-hatching, the late stage of development, was anisotropic, corresponding to the direction of the furrows seen in SEM (Fig. 3c). On the other hand, no such anisotropy was observed in the cell shape of telson primordia at 15 h post-hatching (Fig. 3c). These results suggest that cells acquire anisotropy during development, which determines the direction of furrows. Based on these results, it appears that cuticular furrows occur intercellularly. This study suggests the existence of furrows at various scales, including the multicellular furrows in the horns of *Trypoxylus* (1, 5) and the intracellular furrows in *Drosophila* larvae (34). Each is expected to have a different formation mechanism and should be studied independently.

**Fig. 3.**
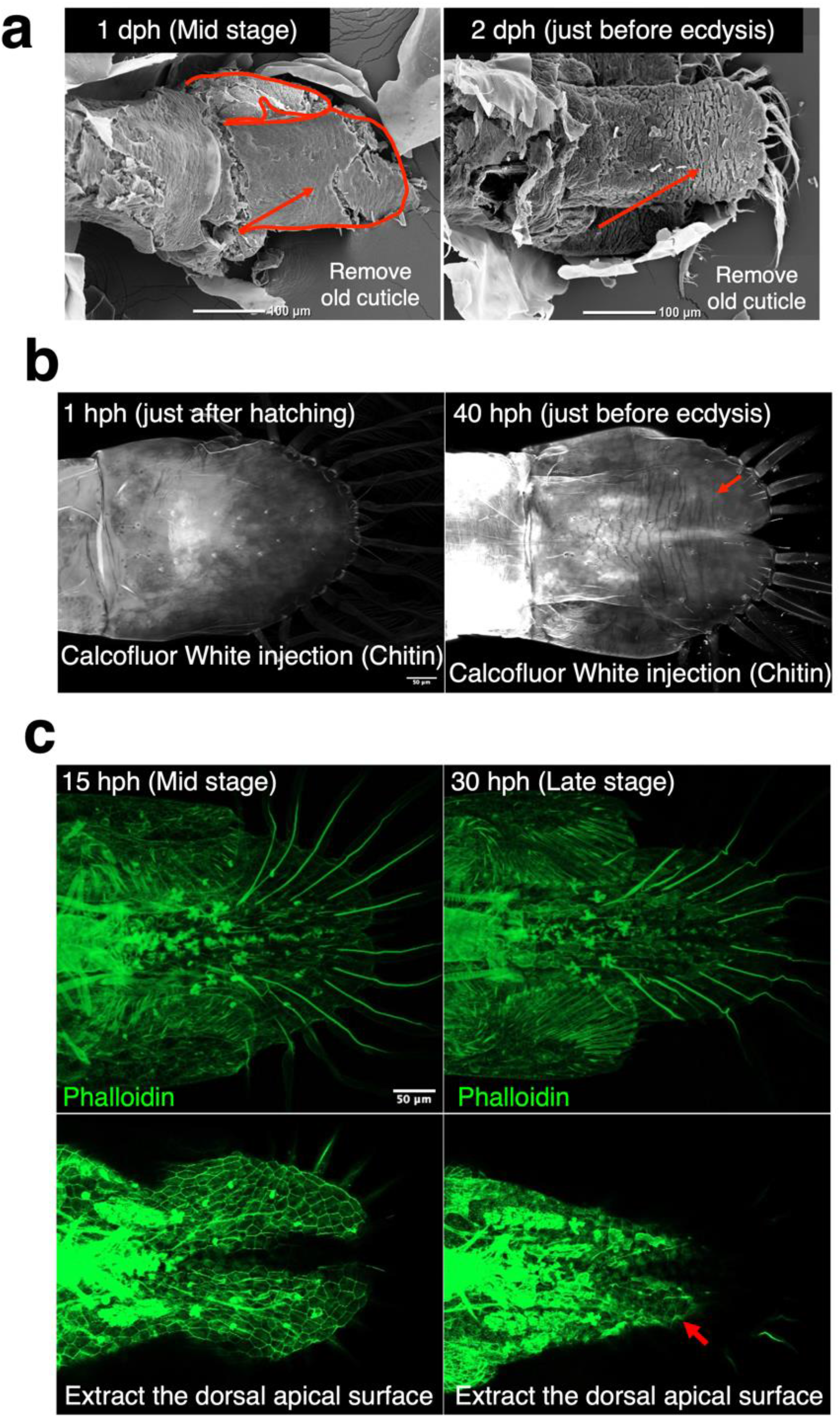
Surface structure of the developmental tail. (a) Scanning electron microscopy (SEM) image of the developmental tail at 1 dph and 2 dph. The red outline shows the uropod and telson branches. The red arrows indicate anisotropic cellular-like shapes (left: mid-stage) and anisotropic cuticular micro furrows (right: just before ecdysis). (b) Calcofluor white injection image of developmental tail at 1hph and 40 hph. The red arrow indicates anisotropic cuticular micro furrows. (c) Phalloidin staining of developmental tail at 15 hph and 40 hph. The top image shows the complete projection, while the bottom image displays the dorsal apical planes projection. The red arrows indicate anisotropic cellular shapes.

### Establishment of *in vivo* live imaging using lectin probe

Next, we thought that the transparent and planar structure of the *Neocaridina* tail would facilitate live observation of the developmental process. In previous research, it was reported that an amazing *in vivo* live imaging system of *Parhyale* using surgery glue could visualize the leg regeneration process up to ecdysis (21). We applied this experimental system to establish an *in vivo* live imaging system using UV-LED resin (Fig. 4a). The use of UV-LED resin significantly reduces the cost compared to the adhesive system used above. On the other hand, UV-LED resins generate heat during curing, so the effect of artefacts due to this must be carefully considered. In the case of *Neocaridina*, the small size of the UV-LED resin allowed them to survive and continue molting, although the time it takes from hatching to ecdysis has slightly increased. The UV-LED resin was placed on either side of the caudal side of the abdomen and the individuals were able to perform abdominal exercises. Some individuals survived ecdysis in live imaging, in which case the tail morphology was normal (Fig. 4b). Our system using fluorescent dye-conjugated WGA-lectin cell labelling has allowed long-term live imaging at 20-minute intervals of the organism and has been able to capture some of the cellular dynamics of the tail from the moment of hatching to the time of ecdysis (Fig. 4b, Mov. 5, 6). WGA allows the visualization of vessels, neurons and hemocytes. In addition, by increasing the exposure, the contours of the tissue can be captured to some extent (Fig. 4b). Observation of the time series of midline optical cross-sections showed that the thickness of the tissue itself did not change much (Fig. 4c). As an optical section, the absolute thickness quantification is not very meaningful, but the relative thickness change is a fact. The fact that the thickness does not change much despite the circulation of hemocytes may suggest to the presence of linkages by dorsal-ventral intercellular projections, as seen in the *Drosophila* wing (35). There are eight setae on the telson of *Neocaridina*. The morphological changes in the telson were analyzed based on the coordinates of the roots of the setae and tips of the telson (Fig. 4d, Mov. 7, 8). In agreement with the results of the fixed tissue analysis (Fig. 1d), it was observed that the width of the telson narrows during development, particularly in the proximal site (Fig. 4e). It was found that hemocytes circulate even during the contraction process of the telson (Mov. 5, 6), suggesting the presence of contractile force generation beyond pressure due to circulation of hemocytes. Tissue shrinkage occurs in various species and scales. Research has shown that morphogenesis in *Drosophila* imaginal discs and beetle horns is influenced by contraction and adhesion (36, 37). Additionally, convergent extension occurs in the embryonic development of various animal species, demonstrating two different mechanistic models of collective cell movements (38). In the future, it is important to consider comparisons with the molecular mechanisms of such phenomena.

**Fig. 4.**
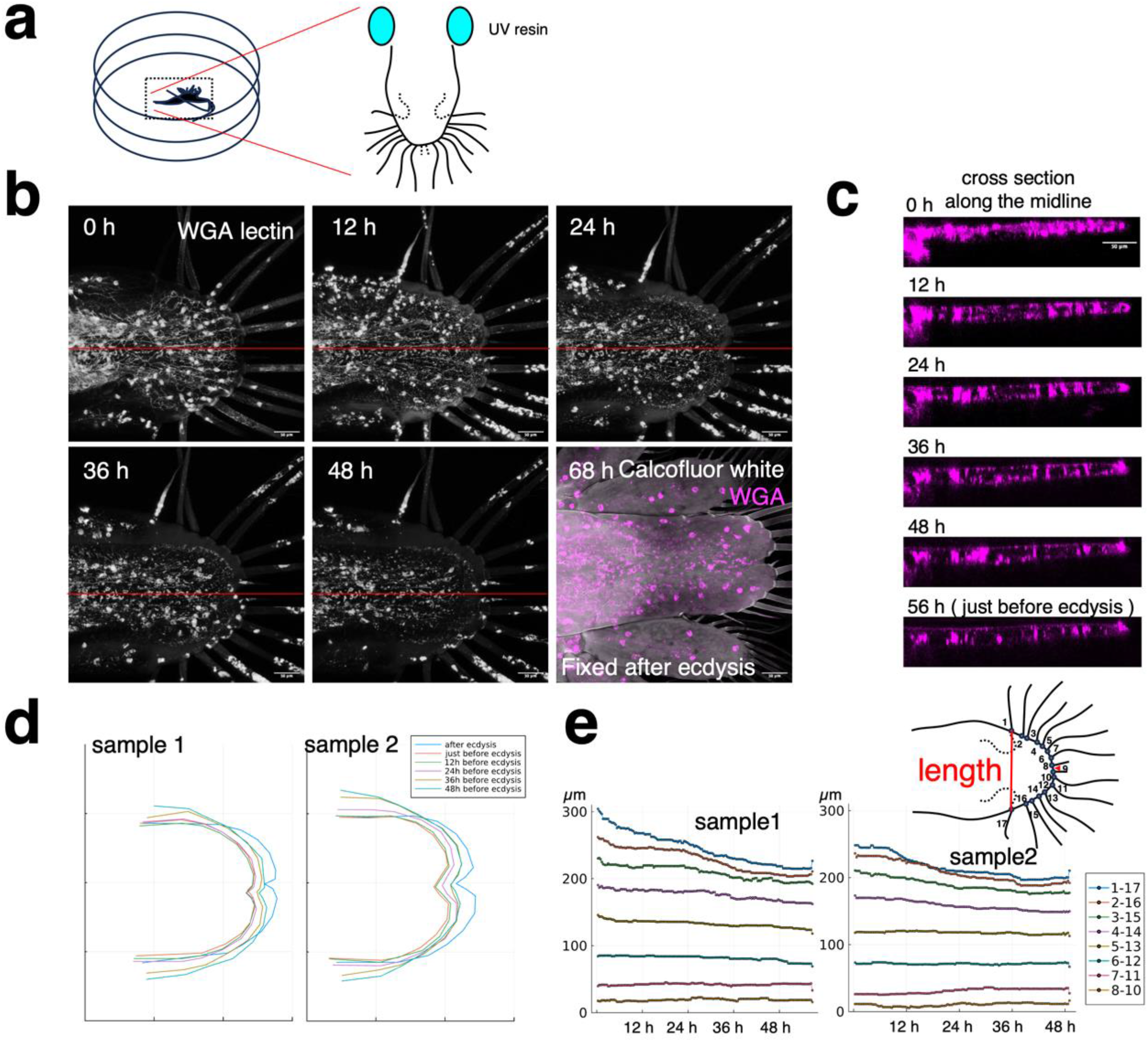
*in vivo* live imaging of the developmental process of the tail. (a) Scheme of immobilization method of a living organism. The UV-LED resin was placed on either side of the caudal side of the abdomen in a glass bottom dish. (b) Time-lapse image of *in vivo* live imaging of the developmental process of the tail. Fluorescent dye-conjugated WGA-lectin injection stains vessels, neurons and hemocytes, the contours of the tissue. 68 hours after starting imaging, the samples after ecdysis were fixed by 4% Paraformaldehyde and stained by Calcofluor white. The images were created by (c) Cross-sectional images of the time-lapse images of the red line (b). (d) The plot of the coordinates of the roots and tips of the setae at some time points of 2 samples. (e) Time series analysis of the width length of the developmental tail. The 8 interval lengths between the left and right setae were plotted.

We noticed that before ecdysis there were hemocyte like cells between the old cuticle and the epithelial tissue. These cells frequently extended their pseudopodia and moved actively during development. They also swelled, burst and died during ecdysis (Movie. 5, Fig. S1). It is thought that ecdysis allowed fresh water to enter from the outside, and the osmotic pressure caused the cells to rupture. Some fluorescence staining images also showed cells outside the epithelial tissue. Apolysis is known to occur in the molting process of conventional arthropods and the gaps are filled with molting fluid, but we are not aware of any reported cases where cells are present. Previous studies have mainly used fixed tissues for analysis, so they may have been washed away in the process. However, in the *Parhyale* leg regeneration experiment, motile cells, probably phagocytes, were reported as “data not shown” in the space between the exoskeleton and the epidermis (21). It is possible that there was an injury as artefact of live imaging in our experiment. However, at least the space between the old cuticle and the epithelial tissue was found to be a viable environment for the cells. Additionally, the previous cuticle, which has shed its epithelium through apolysis and seems to be decomposing, may possess a sturdy framework that inhibits freshwater from penetrating the interior until just before ecdysis.

### Whole genome sequence for expansion of genetic tool

The above analysis shows the developmental process of the tail of *Neocaridina*. This also indicates that *Neocaridina* can be used for *in vivo* live imaging of cell dynamics. However, this study only labeled certain cell types. To monitor the dynamics of a specific cell type, fluorescent transgenic individuals are often generated in model organisms. However, it should be noted that *Neocaridina* currently has only short-read sequence information or unpublished genome assemblies (30, 39, 40). To address this issue, we present a long-read genome sequence of *Neocaridina denticulata* in this study. By utilizing only the nanopore sequence, we were able to capture a draft genome of 2.73 Gb with 36,889 contigs. The longest sequence length were 933 kb. The N50 sequence length was 123 kb and the GC content was 35%. The BUSCO analysis revealed that approximately 80% of the genes in the arthropods database were complete (Table. 1). This sequences can be utilized in the future for various genetic tools, such as fluorescent transgenic lines and detailed live imaging.

**Table 1.**
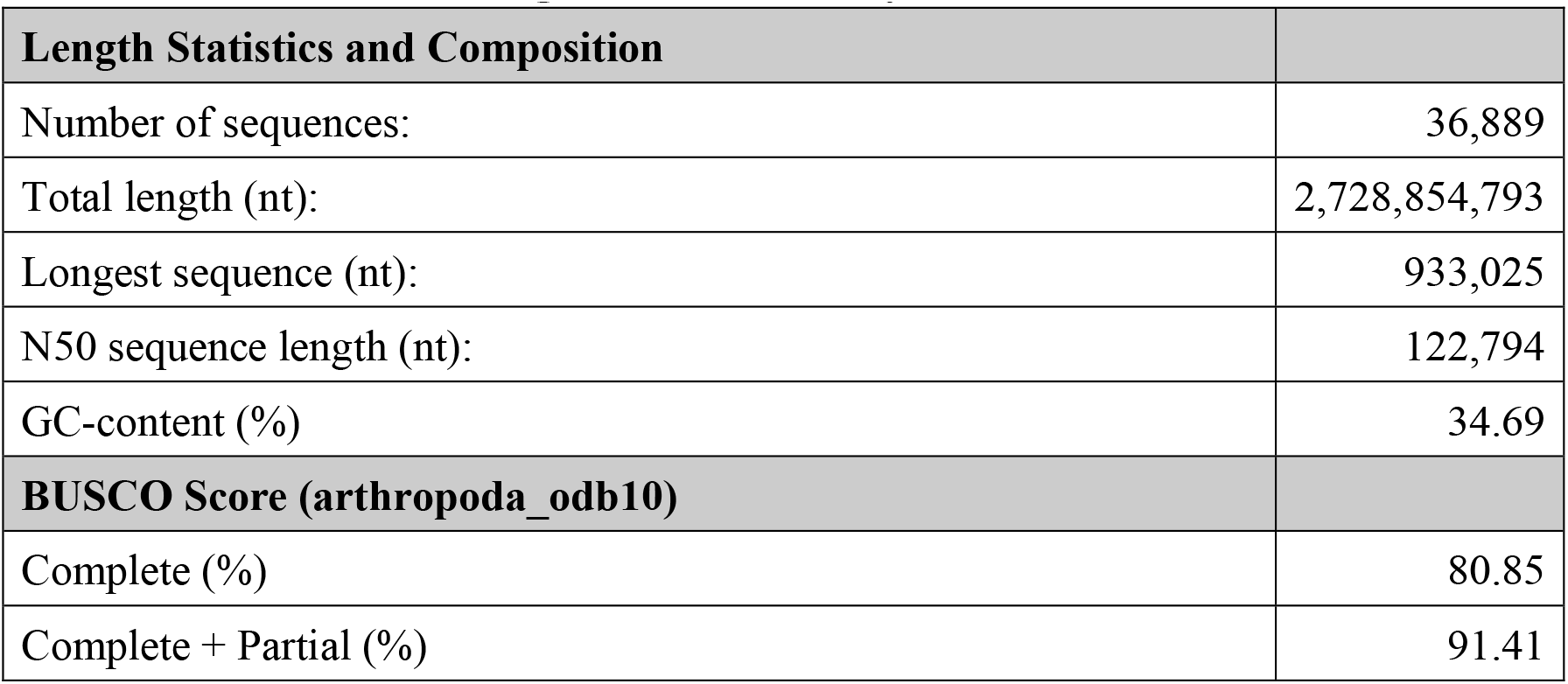
Statistics of the genome assembly.

## Conclusion

Analysis of morphological changes in the tail showed that two major phenomena occur before and after the first ecdysis in *Neocaridina denticulata*. One is a clear branching of the uropod and telson, and the other is a shrinkage and elongation of the telson. Cross-sectional analysis revealed that uropod and telson branching occurs immediately after hatching in the form of cuticle branching. The surface structure of the developmental tail suggests that telson elongation is achieved by the extension of anisotropic furrows in the cuticle during ecdysis. Anisotropy of cuticle furrows is associated with cell shape, and anisotropy of cell shape is found to occur during development from post-hatching. *in vivo* live imaging analysis shows that telson contraction occurs more proximal and gradually prior to ecdysis. These results also demonstrate that *Neocaridina* is useful for cellular and developmental analysis. This study also generated a draft genome of *Neocaridina*, which may stimulate future cell and developmental biology studies with this species.

## Material and Method

### Animals

*Neocaridina denticulata* were commercially purchased and kept in 45 L water tanks at 23-25°C. The tanks were checked daily to identify individuals carrying eggs, which were then transferred to separate bottles. Day 0 was defined as the first day the eggs hatched. Individuals used in time series analysis were checked for hatching every three hours. The hatched individuals were transferred to 96-well plates along with 300 µL of water. The 1st stage larvae mostly underwent ecdysis 40-50 hours after hatching. Movies of the molting were recorded with a BZ-X710 (Keyence, Japan). The captured individuals molted faster, likely due to the higher temperature inside the microscope. The experiments were performed after anesthesia with clove oil diluted approximately 10,000 times.

### Scanning electron microscope (SEM)

The samples were fixed with 4% paraformaldehyde (PFA) in PBS (-) for 24 hours at 4°C and dehydrated using a series of ethanol (50-100%). After dehydration, they were soaked in t-butanol and dried using a freeze-dry system (FZ-2.5; Asahi Life Science, Japan). For observation of the tail primordium, the old cuticle was removed by dissection, using tweezers. Finally, Au sputtering was conducted on the samples with DII-29010SCTR Smart Coater (JEOL Ltd., Japan) and they were observed using a scanning electron microscope JCM-6000plus (JEOL, Japan) with 10 kV.

### Transmission electron microscopy (TEM)

The tails of the samples were isolated and fixed in a solution of 2% paraformaldehyde and 2% glutaraldehyde in PBS (-) at 4°C overnight, before being replaced with PBS. The samples were fixed in 1% osmium tetroxide/0.1 M phosphate buffer at 4°C for 2 hours. They were then dehydrated using a series of ethanol (50-100%) and replaced with QY-1. The samples were then replaced with resin and embedded before being polymerized. The samples were sectioned using an ultramicrotome UC7 (Leica, Germany). Semi-sections cut to around 1 µm were stained by 0.5% toluidine blue. The samples were observed by optical microscope. Ultrathin sections, cut to a thickness of 70-80 nm, were stained with Uranyl acetate solution for 15 minutes, followed by Pb staining solution for 10 minutes. The samples were observed by Transmission electron microscope JEM-1400plus (JEOL, Japan) with 100 kV.

### Fluorescence staining

The tails of the samples were isolated and fixed in 4% PFA 4°C. The fixed samples were treated with 0.1% Triton-X 100 in PBS (-) for 10 min and the samples were soaked in staining solution with 0.1 % Tween-20 in PBS (-). SYTO13 (Thermo Fisher Scientific, USA) was used at a dilution concentration of 1:1000, and Alexa Fluor 488 conjugated phalloidin (Cell Signaling Technology, USA) was used at a dilution concentration of 1:100. Calcofluor White (MP Biomedicals, USA) and CF®555 conjugated WGA (Biotium, USA) or Alexa Fluor 647 Conjugated WGA (Thermo Fisher Scientific, USA) were injected into the living body using microinjector BJ-120 (BEX, Japan) and glass capillary needle. Calcofluor White was used at a mass concentration of 5%, and WGA was used at a concentration of 1 mg/ml. Glass capillary needles were prepared by pullar PC-100 (Narishige, Japan). The injection time was set to ‘short’ mode, and the solution was injected 2-3 times into the larva’s abdomen. For live imaging, the injected individuals were immobilized in the center of a glass bottom dish (Matsunami, Japan) using UV-LED resin (Padico, Japan) and 30 seconds of 395 nm light irradiation. Images were captured using confocal laser scanning microscopy LSM 900 with Axio Observer.Z1 (Zeiss, Germany). Plan-Apochromat 20x/0.8 M27 and Plan-Apochromat 10x/0.45 M27, and GaAsP PMT detector were used. For live imaging, time-lapse images were captured at 20-minute intervals and z-stack imaging was performed at 2µm intervals. The images were processed with ImageJ-Fiji version 2.14.0 (41). Projection figure generated by standard deviation projection type.

### Analysis of the morphology of the telson

Images were analyzed with ImageJ-Fiji version 2.14.0 (41). The length of the telson was measured from the root of the uropod to the tip of the telson. The widths of the telson were measured between the second setae on the proximal side. coordinates of the roots and tips of the setae. For the analysis of the time-lapse images, the coordinates of the roots and tips of the setae were obtained using the “Manual Tracking” plugin. Coordinate information was processed by the Julia language version 1.8. The 8-interval lengths between the left and right setae were calculated from the coordinate information.

### Genome sequence and Genome assembly

Genomic-tip 20/G (QIAGEN, Netherlands) was used to extract genomic DNA from three adult individuals according to the manufacturer’s protocol. The eluted DNA was resuspended in 10 mM Tris-HCl, quantified using Qubit Broad Range (BR) dsDNA assay (Life Technologies, USA) and qualified using TapeStation 2200 with Genomic DNA Screen Tape (Agilent Technologies, USA). Sequencing was performed on a P2 Solo instrument (Oxford Nanopore Technologies, UK) using a PromethION R10.4.1 flow cell. The data obtained were base called using Guppy version 6.4.6 with the ‘super accurate’ base calling mode, and the adapters were trimmed. Reads from each of the three sequencing runs underwent filtration to remove those less than 1000 bp each. Combined set of reads also underwent filtration to remove those less than 4000 bp. Four sets of sequences were prepared due to the computer’s memory limitations. Flye version 2.9.1 was used to assemble sequenced reads from each of the three sequencing runs, as well as from a combined set of reads. The combined set of reads was also assembled by raven-assembler version 1.8.3 (42, 43). The five assemblies were combined into a single assembly using quickmerge version 0.3 (44). Subsequently, the assembly was scaffolded three times using ntlink version 1.3.9 and Flye polishing version 2.9.1 (45, 46). Haplotype duplication was removed and the mitochondrial genome was eliminated using purge_haplotigs version 1.1.2 (47). Contamination was also identified and removed using CAT version 5.3 (48). The draft genome was validated using BUSCO v.5 (49) with the core arthropods (Arthropoda_odb10 database) on the gVolante webserver (50). This sequence is being uploaded to DDBJ under BioSample accession number SAMD00755868.

## Acknowledgments

We appreciate Prof. Shigeru Kondo (Osaka University) and Prof. Shizue Ohsawa (Nagoya University) and Prof. Noriko Funayama (Kyoto University) and their laboratory members for providing an environment for research and helpful discussion. We also thank G-language group (Institute for Advanced Biosciences) for helpful supports and discussion. This research was supported in part by MEXT KAKENHI Grant Number 23K14115 (to HA), 22J00193 (to HA) and 22KJ1542 (to HA) and Mishima Kaiun Memorial Foundation (to HA) and research funds from the Yamagata Prefectural Government and Tsuruoka City, Japan. HA was also supported by Grant-in-Aid for JSPS Fellows (PD).

## Author Contributions

Conceptualization: H.A., Investigation: H.A., Formal analysis: H.A., Funding acquisition: H.A. and K.A. Methodology: H.A. Resources: H.A. and K.A., TEM analysis: N.M. and T.S., Setting up genome sequence: K.A., Writing—original draft: H.A., Writing—review and editing: all authors

---

**Fig. S1.**
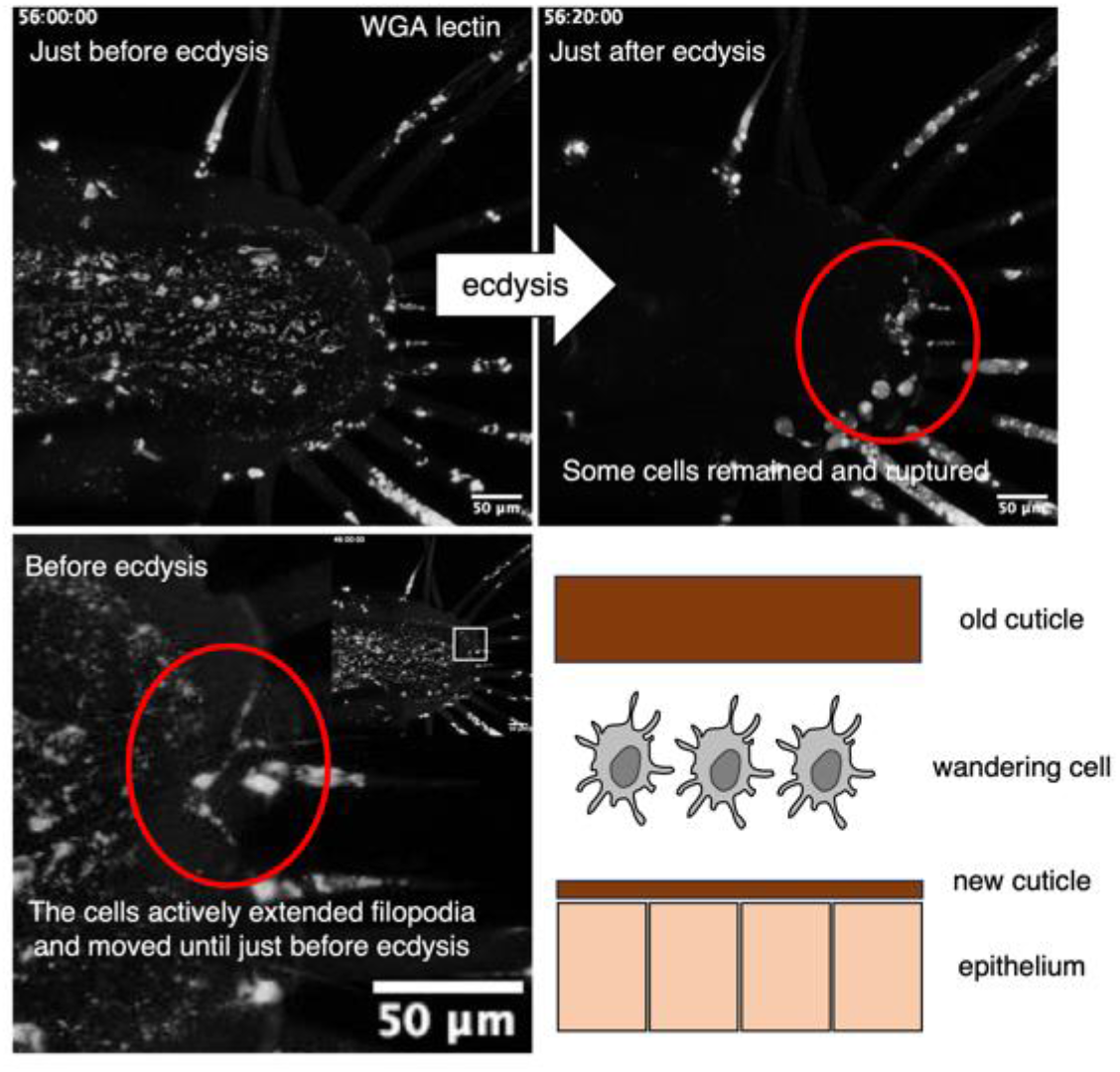
The presence of cells between the old cuticle and epithelial tissue. Time-lapse images just before and after ecdysis. Some cells remained and ruptured after ecdysis, and the cells actively extended filipodia and moved until just before ecdysis.

**Mov. 1 Early stage of 1**^**st**^ **instar development *Neocaridina denticulate* in 96 well plate**

Same sample as Mov. 2. ([link](https://drive.google.com/file/d/1rOP82eql9h3sVr8ch9U7iSy4C2jynPt1/view))

**Mov. 2 Late stage and ecdysis of 1**^**st**^ **instar *Neocaridina denticulate* in 96 well plate**

Same sample as Mov. 1. ([link](https://drive.google.com/file/d/1g3z3pbvnYfzI_MGLm2_DgkkeSoyyHyfG/view))

**Mov.3 z stack imaging of phalloidin staining of 15 hph**

It is played back from the dorsal side to the ventral side. ([link](https://drive.google.com/file/d/1kNrgb2-zABkXfWMqgID8xcQrcXa1C0N2/view))

**Mov.4 z stack imaging of phalloidin staining of 30 hph**

It is played back from the dorsal side to the ventral side. ([link](https://drive.google.com/file/d/1Tqtl3BKldpGhcyULPrvv5xTrTbrrecOn/view))

**Mov. 5 *in vivo* live imaging of the tail development_sample1**

CF®555 conjugated WGA was injected into ∼ 3 hph larva. ([link](https://drive.google.com/file/d/1lBxRjGKxXNj1xx77_53tG1iU9_DbTEkv/view))

**Mov. 6 *in vivo* live imaging of the tail development_sample2**

Alexa Fluor 647 conjugated WGA was injected into around 10 hph larva. ([link](https://drive.google.com/file/d/1ZhL-UQT9TIHsLakCMxU6AynQInaDidBX/view))

**Mov. 7 Morphological changes in the telson_sample1**

Manual tracking of the coordinates of the roots and tips of the setae. ([link](https://drive.google.com/file/d/1ocMr45CJY02YBqct0EM69B3A6q33LMHo/view))

**Mov. 8 Morphological changes in the telson_sample2**

Manual tracking of the coordinates of the roots and tips of the setae. ([link](https://drive.google.com/file/d/1ojRz4fioQ73RZWRY74EVvZzFNLYGnEDk/view))

